# Imp1 acts as a dosage- and stage-dependent temporal rheostat orchestrating radial glial fate transitions and cortical morphogenesis

**DOI:** 10.1101/2025.11.18.688993

**Authors:** Romie Angelo G. Azur, Daniel Feliciano, Isabel Espinosa-Medina, Raghabendra Adhikari, Joaquin Lilao-Garzón, Ella Jensen, Ching-Po Yang, Tzumin Lee

## Abstract

Cortical neurogenesis proceeds through a precise temporal program in which radial glia sequentially generate distinct neuronal subtypes and later glia, yet how post-transcriptional regulators coordinate these transitions remain poorly understood. We previously identified that a decreasing temporal gradient of the RNA-binding protein Imp encodes neural stem cell age in *Drosophila*. In this work, we extend our investigation to Imp1, a mammalian homologue of Imp, and its role in murine neocortical development. Using TEMPO to track birth-order dynamics, we demonstrate that sustained Imp1 overexpression during early neurogenesis arrests temporal fate progression, shifting neuronal populations toward deeper cortical layers V-VI. Immunostaining with layer-specific transcription factors Cux1 and Ctip2 confirmed that laminar repositioning results from genuine changes in neuronal identity rather than migratory defects, with neurons adopting molecular identities matching their final positions. Temporal window-specific manipulations reveal distinct stage-specific effects where early-stage Imp1 induction produces cascading effects on fate specification and moderately delays the neuronal-to-gliogenic transition, while mid-stage induction induces neuronal accumulation in the subplate region. Live imaging of organotypic cultures reveals continuous neuronal recruitment within intermediate and ventricular zones, with mid-stage-born neurons accumulating at significantly faster rates than earlier cohorts. Strikingly, mid-stage Imp1 overexpression also induces ectopic glial-like foci distributed throughout the cortical plate, featuring dramatic cellular expansion and morphological heterogeneity. These findings establish Imp1 as a dosage- and stage-dependent temporal rheostat orchestrating developmental transitions in radial glial progenitors, controlling neuronal fate decisions and spatial organization. This work advances our understanding of molecular timing mechanisms governing neuronal diversity in the mammalian cortex.

## INTRODUCTION

The formation of the mammalian neocortex relies on precise temporal patterning mechanisms that govern when distinct neuronal subtypes are generated during development. Cortical projection neurons are born in a stereotyped inside-out sequence, with early-born neurons (E11-E13) populating deep layers VI and V, while later born neurons (E14-E16) migrate past their predecessors to occupy superficial layers II-IV (Angevine & Sidman, 1961; Leone et al., 2008; Molyneaux et al., 2007; Rakic, 1974). This inside-out organization is fundamental to establishing proper cortical circuitry and function (Cooper, 2008; Gadisseux et al., 1990; S. He et al., 2015; Rakic, 1972, 1988). Understanding how this precise temporal organization emerges during development requires examining the regulatory mechanisms that control when different neuronal types are generated.

Recent advances have revealed that this temporal precision emerges through coordinated control systems operating at multiple regulatory levels. At the transcriptional level, single-cell profiling has revealed that RGCs display dynamic gene expression over developmental time, with early RGCs expressing higher levels of genes like Hes5 and late RGCs showing elevated expression of factors such as Pou3f2 (Ruan et al., 2021). At the epigenetic level, radial glial cells (RGCs) progress through temporal “ground states” that are heritably transmitted through their progeny, with apical progenitors retaining remarkable plasticity to revert to earlier competence states when exposed to younger cellular environments (Oberst et al., 2019; Telley et al., 2019). Metabolic mechanisms provide an additional temporal control layer, as forebrain-enriched proteins like FAM210B sustain extended neurogenesis by lengthening cell cycles and reducing consumptive divisions compared to more transient hindbrain neurogenesis (Baumann et al., 2025).

These multilayered temporal control systems coordinate two distinct neurogenic modes that generate cortical complexity: direct neurogenesis produces the major projection neuron classes while indirect neurogenesis creates specialized subtypes within each class to assemble fine-scale cellular mosaics (Huilgol et al., 2023). Importantly, the resulting lamination patterns prove more nuanced than the classical inside-out model suggests, neurons destined for different layers can be born simultaneously, same-layer neurons may arise at different developmental times, and simultaneously-born neurons exhibit progressively decreasing fate heterogeneity as development advances (Huilgol et al., 2025; Magrinelli et al., 2022).

While these studies have illuminated transcriptional and metabolic mechanisms, the contribution of post-transcriptional regulatory layers to temporal precision in cortical development remains largely unexplored. This raises the question of whether additional regulatory layers, particularly post-transcriptional mechanisms, contribute to temporal precision in cortical development.

RNA-binding proteins that post-transcriptionally regulate gene expression programs represent key candidates for filling this knowledge gap (Cesari et al., 2024; Lennox et al., 2018; Pilaz & Silver, 2015). Among these, the Imp (IGF2 mRNA-binding protein) family has emerged as a critical regulator of temporal identity in diverse developmental contexts (Degrauwe et al., 2016; Lee et al., 2025; X. Li et al., 2025; Yaniv & Yisraeli, 2002). In *Drosophila*, Imp exhibits a characteristic temporal gradient during neuroblast divisions with high early expression promoting early temporal fates and gradual decline allowing progression to later fates (Liu et al., 2015). This temporal gradient appears evolutionarily conserved, as mammalian Imp1 (also known as Igf2bp1 and Zbp1) shows similar expression dynamics in cortical progenitors (Nishino et al., 2013). Loss-of-function studies confirm Imp1’s developmental importance as complete knockout exhibits perinatal lethality with disrupted cortical organization, impaired neuronal migration and reduced progenitor activity during the E12.5 – E17.5 neurogenic period (Hansen et al., 2004; Núñez et al., 2022), while partial loss accelerates premature gliogenesis (Nishino et al., 2013). These findings establish Imp1 as essential for proper temporal patterning in mammalian corticogenesis.

Despite compelling evidence for Imp’s importance in temporal fate specification, functional studies in the mammalian cortex are often constrained by the lack of tools to manipulate and visualize temporal cell fate decisions with sufficient resolution. Traditional approaches using constitutive genetic perturbations often obscure stage-specific effects by altering gene expression throughout development (Bolt & Duboule, 2020; Hippenmeyer, 2023), while conventional lineage tracing methods typically cannot distinguish between consecutive cell generations born from the same progenitor pool (Hsu, 2015; Mao et al., 2025; Richier & Salecker, 2015). These constraints have prevented detailed analysis of how temporal regulators modulate specific developmental transitions, making it difficult to determine when and how these factors exert their effects.

The recent development of TEMPO (Temporal Encoding and Manipulation in a Predefined Order) has provided a solution to these challenges (Espinosa-Medina et al., 2023). This CRISPR-based system enables simultaneous visualization of sequentially-born cell cohorts through distinct fluorescence reporters and temporal manipulation of gene expression within defined developmental windows. Unlike previous approaches, TEMPO allows researchers to track developmental history of cells while manipulating molecular constituents with precise temporal control. TEMPO delivery through *in utero* electroporation creates mosaic perturbations by targeting a select RGC cohorts within an otherwise normal developmental environment. This approach is particularly well-suited for dissecting cell-autonomous and intrinsic temporal fate regulatory mechanisms.

Here, we employ TEMPO to systematically investigate how Imp1 regulates temporal fate progression in the developing mouse neocortex. Through continuous and temporally-restricted Imp1 overexpression paradigms, we demonstrate that Imp1 functions as a critical temporal fate regulator that arrests normal developmental progression when ectopically maintained. Our findings reveal that Imp1 exhibits remarkable sensitivity to expression timing, with distinct temporal windows exhibiting differential responses to perturbation. Sustained Imp1 expression promotes retention of early neuronal identities, shifting neuronal populations toward deep layer fates while suppressing the normal transition to gliogenesis. We uncover temporal window-specific organizational defects and abnormal glial accumulation phenotypes. Early Imp1 activity produces cascading effects on neuronal specification that extend into subsequent temporal cohorts and moderately delays gliogenesis, whereas mid-stage expression induces subplate neuronal accumulations and dramatic glial foci formation without affecting the neurogenic-to-gliogenic transition. These findings establish Imp1 as a temporally sensitive developmental rheostat that coordinates both cell fate decisions and tissue organization in the mammalian cortex.

## MATERIALS AND METHODS

### Experimental Model

All mouse experiments were conducted in accordance with NIH guidelines for animal research and approved by the Institutional Animal Care and Use Committee at Janelia Research Campus, Howard Hughes Medical Institute. Mice were maintained at 28°C under a 12h light/dark cycle in temperature-controlled rooms. Experiments utilized embryos (E12.5-E16.5) and postnatal day 10 (P10) pups from timed-pregnant C57BL/6J females (Charles River Laboratories). Both male and female mice were used for postnatal analyses.

### *In utero* Electroporation and TEMPO Induction

TEMPO visualization during cortical development was achieved through *in utero* electroporation performed at embryonic days E12.5, E13.5 or E14.5 as previously described (Loulier et al., 2009; Wang & Mei, 2013). Pregnant females were anesthetized with 2% isofluorane in oxygen and uterine horns were exposed via midline incision. DNA solution (1 µL) containing equimolar concentrations of interdependent TRE:TEMPO-gRNA-1trigger, CAG:Cas9-2A-tTA-gRNA-2switch and CAG: PiggyBac Transposase plasmids (total DNA concentration: 2 µg/µL) supplemented with Fast Green dye was pressure-injected into the lateral ventricle using pulled glass capillaries. Electroporation was performed using tweezer electrodes with four 50 ms pulses at 100 V delivered at 950 ms intervals. Following surgical closure, animals were allowed to recover in clean cages before collection at embryonic stages (E16.5) or birth for postnatal analysis.

### Tissue Processing and Immunohistochemistry

P10 mice were deeply anesthetized with 5% isofluorane and transcardially perfused with cold saline followed by 4% PFA. Brains were post-fixed overnight in 4% PFA at 4°C, then sectioned at 100 µm. Immunostaining was performed in blocking buffer (PBS containing 2% bovine serum albumin and 0.8% Triton X-100). Primary antibodies were applied overnight to amplify TEMPO signals or detect fate markers at 4°C: mouse anti-V5 (1:650, Thermo Fisher R96025) to detect CFP+ cells, rat anti-mCherry (1:500, Thermo Fisher M11217) or chicken anti-mCherry (1:500, AvesLabs MCHERRY AB_2910557) to detect RFP+ cells, rat anti-Ctip2 (1:100, Abcam [25B6] ab18465) to detect Ctip2+ cells, and rabbit anti-Cux1 (1:50, Proteintech 11733-1-AP) to detect Cux1+ cells. Following three 15-minute washes, sections were incubated with secondary antibodies for 2-4 hours at room temperature: Alexa Fluor 647 goat anti-mouse (1:500, Thermo Fisher A-21235) and Alexa Fluor 568 goat anti-rat (1:500, Thermo Fisher A-11077). Processed sections were mounted using Fluoromount-G and imaged using Zeiss LSM 880 and Leica STERLLARIS confocal microscopes. Image analysis was performed using Fiji for cell quantification and morphological assessments.

### Live cell imaging

To monitor cell migration of TEMPO+ cells in real-time, we adapted established organotypic slice culture protocols with modifications for extended imaging (Wiegreffe et al., 2017). Briefly, TRE:TEMPO-gRNA-1trigger and CAG:Cas9-2A-tTA-gRNA-2switch plasmids were electroporated into E12.5 embryonic cortices, to be harvested 48 hours later. Brains were rapidly dissected in ice-cold complete HBSS solution (50 mL 10X HBSS, 1.25 mL 1M HEPES buffer pH 7.4, 15 mL 1M D-glucose, 5 mL 100 mM calcium chloride, 5 mL 100 mM magnesium sulfate, and 2 mL 1M sodium bicarbonate in 500 mL sterile water). Dissected tissue was embedded in 3% low melting point agarose (Invitrogen, 16520100) maintained at 38°C and sectioned at 250 µm thickness using a vibratome (Leica VT1200S) with parameters optimized for embryonic tissue: 0.99 mm amplitude and 0.09-0.12 mm/s speed. Sections were transferred to 6-well plates containing ice-cold complete HBSS and screened under fluorescence stereomicroscopy to identify slices with robust CFP expression in cortical regions. Selected brain slices were placed on cell culture membrane inserts (Millipore PICM0RG50) in 6-well plates containing slice culture medium (35 mL Basal Medium Eagle, 12.9 mL complete HBSS, 1.35 mL 1M D-glucose, 0.25 mL 200 mM L-glutamine, 0.5 mL penicillin-streptomycin, supplemented with 5% horse serum, final volume 50 mL, sterile filtered). Time-lapse acquisition was performed using an inverted Zeiss LSM 980 confocal microscope equipped with a 20X dry objective and environmental chamber maintaining 37°C and 5% CO_2_. Slice preparations were transferred to 50 mm glass-bottom dishes containing 2 mL culture medium for imaging. Acquisition parameters included 2.5 µm z-steps captured every 30 minutes over 48 hours total duration. Time-lapse datasets were analyzed using Fiji and Imaris software to track individual cell behaviors and quantify TEMPO color transitions during cortical development.

### Quantitative Analysis and Statistical Methods

Following confocal imaging, quantitative analysis of TEMPO reporter distribution was performed using automated cell detection algorithms to ensure objective measurements and minimize manual bias in cell counting. The total number of TEMPO-positive cells expressing each fluorescent reporter (CFP, RFP, YFP) was determined using a custom FIJI macro incorporating StarDist neural network-based cell detection and segmentation (Schmidt et al., 2018; Weigert & Schmidt, 2022). Prior to automated analysis, regions of interest were carefully defined based on anatomical landmarks visible through DAPI nuclear counterstaining. Cortical layers were delineated into upper layers (II-IV), lower layers (V-VI) and the subplate zone (SPZ).

For quantification of subplate zone and glial populations, image samples were first blinded to experimental conditions and individual foci were manually counted using FIJI’s built-in cell counter tool to complement the automated analysis pipeline. The same manual approach was used to quantify cells co-expressing TEMPO reporters alongside established cortical layer markers.

Given the inherent variability in electroporation efficiency across individual embryos, raw cell counts were normalized to account for differences in overall labeling density. Data were presented as the percentage of neurons distributed within upper versus lower cortical layers or the proportion of glial cells, for each TEMPO color reporter. Additionally, we calculated the relative percentage of each TEMPO reporter within the total TEMPO-positive population to control for potential alterations in reporter cascade progression that might confound interpretation of experimental manipulations. Statistical comparisons between control and experimentally perturbed conditions were performed using two-tailed unpaired Welch’s t-test in GraphPad Prism software.

## RESULTS

### Sustained ectopic Imp1 expression during early neurogenesis alters cortical laminar organization and promotes cell accumulation at the subplate

To investigate how Imp1 regulates temporal fate specification during cortical neurogenesis, we first examined the effects of continuous Imp1 overexpression throughout cortical development. We electroporated constructs for constitutive Imp1 expression into mouse embryonic cortices at either E12.5 or E13.5 using our TEMPO system, which sequentially labels temporal windows of neuronal birth through color transitions occurring approximately every 24 hours, to simultaneously track birth-order and laminar position (Figure 1A) (Espinosa-Medina et al., 2023). Following E12.5 electroporation, T1 (CFP+) marks neurons born until E13.5, T2 (RFP+) corresponds to E13.5-E14.5, and T3 (YFP+) represents E14.5-E15.5 and later stages, enabling precise temporal resolution of fate decisions across cortical development.

**Figure 1.**
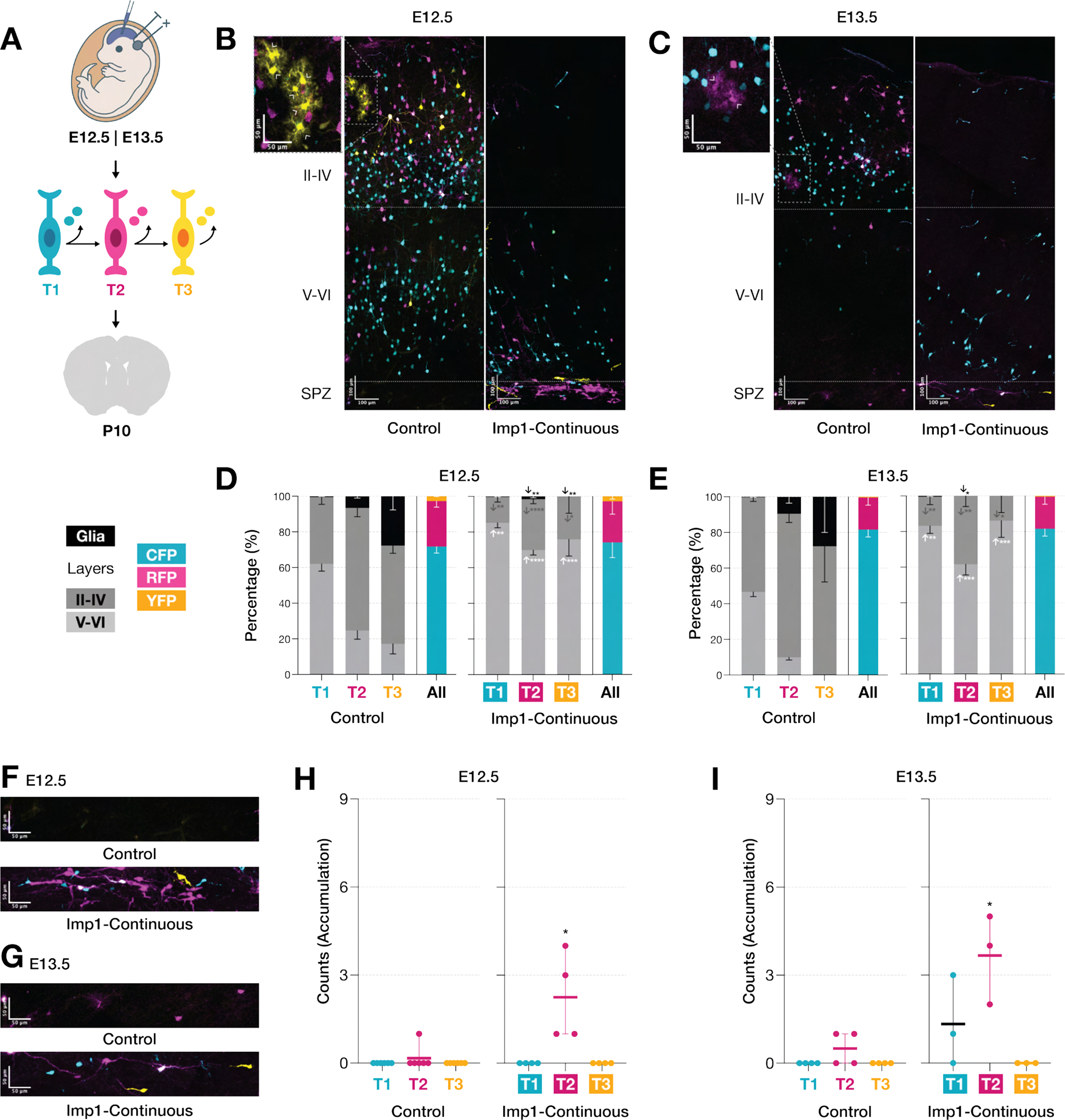
Continuous Imp1 overexpression induced during early neurogenesis redistributes neuronal laminar organization and promotes cell accumulation at the subplate. **(A)** Experimental paradigm: *in utero* electroporation of TEMPO. TEMPO reporters (CFP, RFP, YFP) label sequential temporal windows of neuronal birth (T1, T2, T3). **(B,C)** Representative cortical sections showing TEMPO distribution in control vs continuous Imp1 induction. Insets beside control setups show normal glia clusters (outlined arrowheads) **(D,E)** Quantification reveals disrupted laminar organization with redistribution toward deeper layers (left) while TEMPO proportions remain unchanged (right) suggesting arrested temporal progression. **(F,G)** High magnification images show cellular accumulations beneath layer VI. **(H,I)** Subplate accumulations are enriched for T2-labeled neurons (E12.5) or T1+T2 neurons (E13.5), indicating temporal window-specific sensitivity. Dashed lines: upper (II-IV), lower cortical layers (V-VI) and subplate zone (SPZ). Scale bars represent (B,C) 100 µm and (F,G) 50 µm. Data represent mean±SEM. Statistical significance was calculated by comparing each experimental condition within each temporal population with their corresponding control counterpart using two-tailed unpaired Welch’s t-test (*P < 0.05, **P < 0.01, *** P < 0.001).

Analysis at postnatal day 10 (P10) revealed two striking phenotypes from sustained Imp1 overexpression. The first phenotype involved significant disruption of canonical inside-out layer formation alongside pronounced suppression of gliogenesis (Figure 1B,C). Control electroporations faithfully recapitulated normal developmental patterns with early-born T1 neurons (CFP+) predominantly occupying deep layers V-VI, while later born T2 (RFP+) and T3 (YFP+) neurons populating progressively more superficial positions (Figure 1D). When initiated at E13.5, this laminar pattern shifted superficially as expected from the delayed induction timing (Figure 1E).

Continuous Imp1 overexpression significantly altered this organization, redistributing neurons from all temporal windows toward deeper cortical layers. This effect was particularly pronounced for T2 and T3 populations, which normally contribute predominantly to superficial layer formation but were instead redirected to deeper layers following Imp1 overexpression (Figure 1D,E). Importantly, the relative proportions of each TEMPO color population remained unchanged across conditions (Figure 1D,E, colored panels), indicating that this laminar redistribution more likely reflects altered neuronal fate specification rather than perturbed cell cycle or TEMPO transition kinetics.

Accompanying these laminar changes was a marked suppression of gliogenesis. In control conditions, glial cells emerged around the T2 temporal window and significantly increased in number during T3, reflecting the normal developmental transition from neurogenesis to gliogenesis. Imp1 overexpression dramatically reduced glial cell numbers across multiple temporal windows following E12.5 electroporation (Figure 1D). With E13.5 electroporations, we observed a significant reduction in T2 glia, with T3 showing a similar trend that did not reach statistical significance, possibly due to limited sample size (Figure 1E). These observations align well with previous reports that Imp1 loss-of-function leads to premature astrocyte generation (Nishino et al., 2013), which together with our findings, establishes Imp1 as a key regulatory factor that maintains progenitor cells in a neurogenic state and prevents premature entry into gliogenesis.

The second phenotype involved the unexpected accumulation of TEMPO+ cells within the subplate zone positioned beneath layer VI (Figure 1F,G). These ectopic cellular clusters were particularly enriched in T2-labeled cells following E12.5 electroporations (Figure 1H). When electroporation timing shifted to E13.5, the foci composition remained predominantly composed of T2-labeled cells but also included some T1-labeled neurons (Figure 1I). This shifting composition reveals temporal window-specific sensitivity to Imp1-mediated perturbations, suggesting that different developmental stages possess distinct vulnerabilities to Imp1 manipulation.

The emergence of these two striking phenotypes over days of development, altered laminar organization with suppressed gliogenesis and abnormal subplate accumulation, raised critical questions about the temporal specificity underlying Imp1’s effects. To dissect whether these alterations result from cumulative effects throughout development or reflect specific temporal requirements, we next sought to examine the consequences of restricting Imp1 expression to discrete developmental windows.

### Temporal window-specific Imp1 overexpression reveals differential stage sensitivity on cortical laminar fate and subplate organization

To determine whether Imp1’s effects depend on specific developmental contexts and to identify critical periods for temporal fate regulation, we employed TEMPO constructs that restrict Imp1 expression to individual temporal windows (T1 or T2) following E12.5 electroporation. We also attempted T3-restricted expression but the resulting low numbers of YFP+ neurons precluded any meaningful analysis.

Restricting overexpression to discrete temporal windows produced markedly different effects from continuous expression. While continuous Imp1 overexpression caused dramatic developmental arrest of temporal fate progression, T1-specific induction (T1^Imp1^) allowed partial preservation of laminar organization while still biasing neurons toward deeper positions (Figure 2A). The T1 population showed enhanced retention in layers V-VI compared to controls and this deep-layer bias persisted into the RFP+ T2 cohort (Figure 2B), showing that early Imp1 activity influences subsequent temporal windows beyond the initial induction window. However, in contrast to the near-complete suppression of gliogenesis observed with continuous expression, T1^Imp1^ overexpression produced only moderate reductions in glial cell numbers across temporal cohorts, suggesting that restricting Imp1 induction to early-stage delays but does not block the neurogenesis-to-gliogenesis transition.

**Figure 2.**
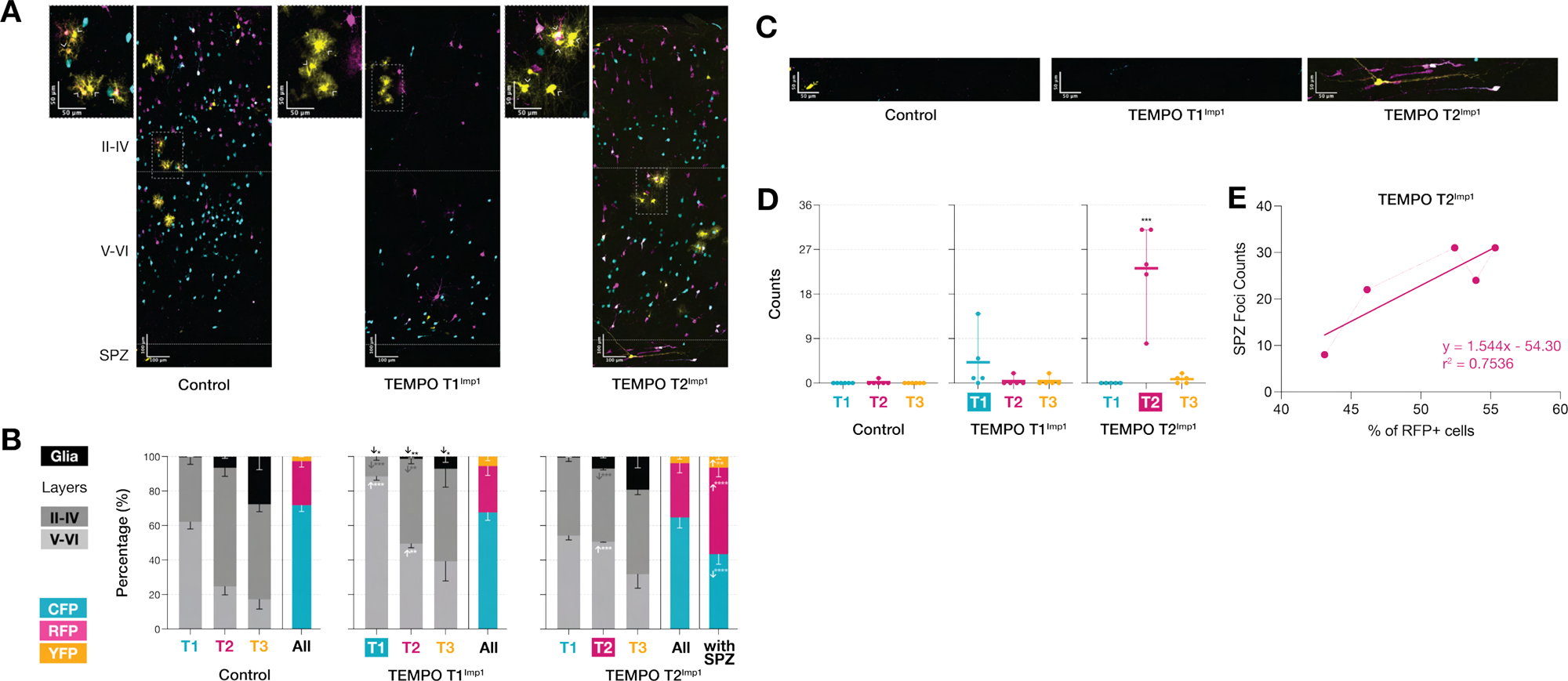
Temporal window-specific Imp1 induction reveals stage-restricted effects of Imp1 on cortical laminar fate and subplate organization. **(A)** Representative images of TEMPO distribution following T1- or T2-restricted Imp1 overexpression via IUE at E12.5. Insets beside each setup show normal glia clusters (outlined arrowheads). **(B)** T1^Imp1^ redistributes neurons towards deeper layers with cascading effects on T2, and reduces glia across cohorts. T2^Imp1^ shows restricted effects on laminar positioning without affecting glia or T3 cohorts. When SPZ is excluded, TEMPO proportions remain unchanged in T2^Imp1^ brains vs controls. Including SPZ foci increased %RFP+ and %YFP+ at the expense of %CFP+. **(C)** High magnification images of accumulations in SPZ following T2^Imp1^. **(D)** Quantification confirms increased SPZ foci only in T2^Imp1^. **(E)** Positive correlation between RFP+ SPZ foci number and %RFP+ cells (r² = 0.7536, slope = 1.544). Dashed lines: upper (II-IV), lower cortical layers (V-VI) and subplate zone (SPZ). Scale bars represent (A) 100 µm and (C) 50 µm. Data show mean±SEM. Statistics: two-tailed unpaired Welch’s t-test (*P < 0.05, **P < 0.01, ***P < 0.001).

T2-restricted Imp1 overexpression (T2^Imp1^) revealed distinct temporal specificity from continuous and T1^Imp1^. As expected, T1 neurons showed no changes in laminar positioning since Imp1 expression was not yet active during their generation. However, T2 neurons were redistributed toward deeper layers without affecting the subsequent T3 cohort, a pattern distinct from both T1^Imp1^ and continuous expression conditions where effects cascaded into later temporal windows. Additionally, T2^Imp1^ overexpression did not reduce the generation of normally-dispersed glial cells, demonstrating temporally restricted effects on the gliogenic transition. Most notably, T2^Imp1^ uniquely induced ectopic accumulations within the subplate zone that were specifically enriched in RFP-labeled (T2) neurons (Figure 2C,D).

To further characterize these SPZ accumulations, we examined the overall distribution of TEMPO populations across the entire brain. Quantification of TEMPO proportions within cortical layers alone (excluding SPZ) reveled statistically unchanged distributions of CFP+, RFP+, and YFP+ populations in T2^Imp1^ brains compared to controls (Figure 2B, left colored bar). However, when SPZ foci were included in the analysis, we observed a significant shift toward increased %RFP+ and %YFP+ cells at the expense of %CFP+ populations (Figure 2B, right colored bar). Furthermore, the number of RFP+ SPZ foci positively correlated with the overall percentage of RFP+ cells when SPZ was included in the quantification (Figure 2E) (r^2^ = 0.7536, slope = 1.544), indicating that brains exhibiting more severe SPZ foci formation showed correspondingly greater representation of RFP+ neurons overall. This finding directly parallels our observations from continuous overexpression, where subplate accumulations following E12.5 electroporations were similarly dominated by RFP+ neurons, indicating that this organizational defect represents a T2-specific response to Imp1 perturbation.

These temporal-window specific manipulations reveal that Imp1’s capacity to regulate cortical development is stage-dependent. Early Imp1 activity (T1) produces broad, cascading effects on both neuronal fate specification and gliogenesis timing, while later expression (T2) shows more restricted influence on laminar positioning but unique sensitivity for inducing subplate organizational defects. The progressive reduction in Imp1’s regulatory scope from the comprehensive developmental arrest seen with continuous expression to the increasingly focused effects of T1 and T2-restricted expression reveals stage-dependent potency. This suggests that Imp1’s temporal fate regulatory mechanisms are more potent during early cortical neurogenesis and become progressively constrained as neuronal precursors transition to subsequent cell states.

### Laminar redistribution via temporal Imp overexpression reflects genuine changes in neuronal cell fate specification

The dramatic shifts in laminar positioning observed following Imp1 overexpression could result from either altered neuronal fate specification or disrupted migration patterns that misplace correctly specified neurons. To distinguish between these possibilities, we examined layer-specific transcription factor expression in TEMPO-labeled neurons by immunostaining P10 brain sections for Cux1 (upper-layer marker) and Ctip2 (deep-layer marker) (Arlotta et al., 2005; Cánovas et al., 2015; García-Moreno & Molnár, 2015; Nieto et al., 2004). We focused our analysis on CFP+ neurons located in deep layers V-VI, as this population represented the largest cohort within the lower layers following Imp1 overexpression and thus provided the most robust dataset for assessing molecular identity changes. We also examined RFP+ neurons in layers V-VI to determine whether later-born neurons similarly adopt appropriate deep-layer identities when redirected to these positions, though the smaller numbers of T2-born neurons normally residing in deep layers necessitate cautious interpretation due to higher variance and limited sampling.

Control conditions first established the expected relationship between neuronal identity and laminar position during normal cortical development (Figure 3A,B). Within the T1 temporal window, approximately 78% of CFP+ cells in lower layers expressed Ctip2, while only 3% expressed Cux1 (Figure 3C, left). The remaining 19% were double-negative for both markers. These double negatives likely represent neurons adopting less canonical layer identities that do not conform to the standard Cux1/Ctip2 dichotomy (Bandler et al., 2022). RFP+ neurons in layers V-VI showed a more distributed marker profile in controls, with roughly equal proportions expressing Ctip2 (50%) or neither marker (47%), and minimal representation of Cux1+ (2%) or double-positive cells (1%) (Figure 3C, right). This distribution suggests that T2-born neurons normally populate primarily superficial layers, with only a subset migrating to or remaining in deep layers during normal development.

**Figure 3.**
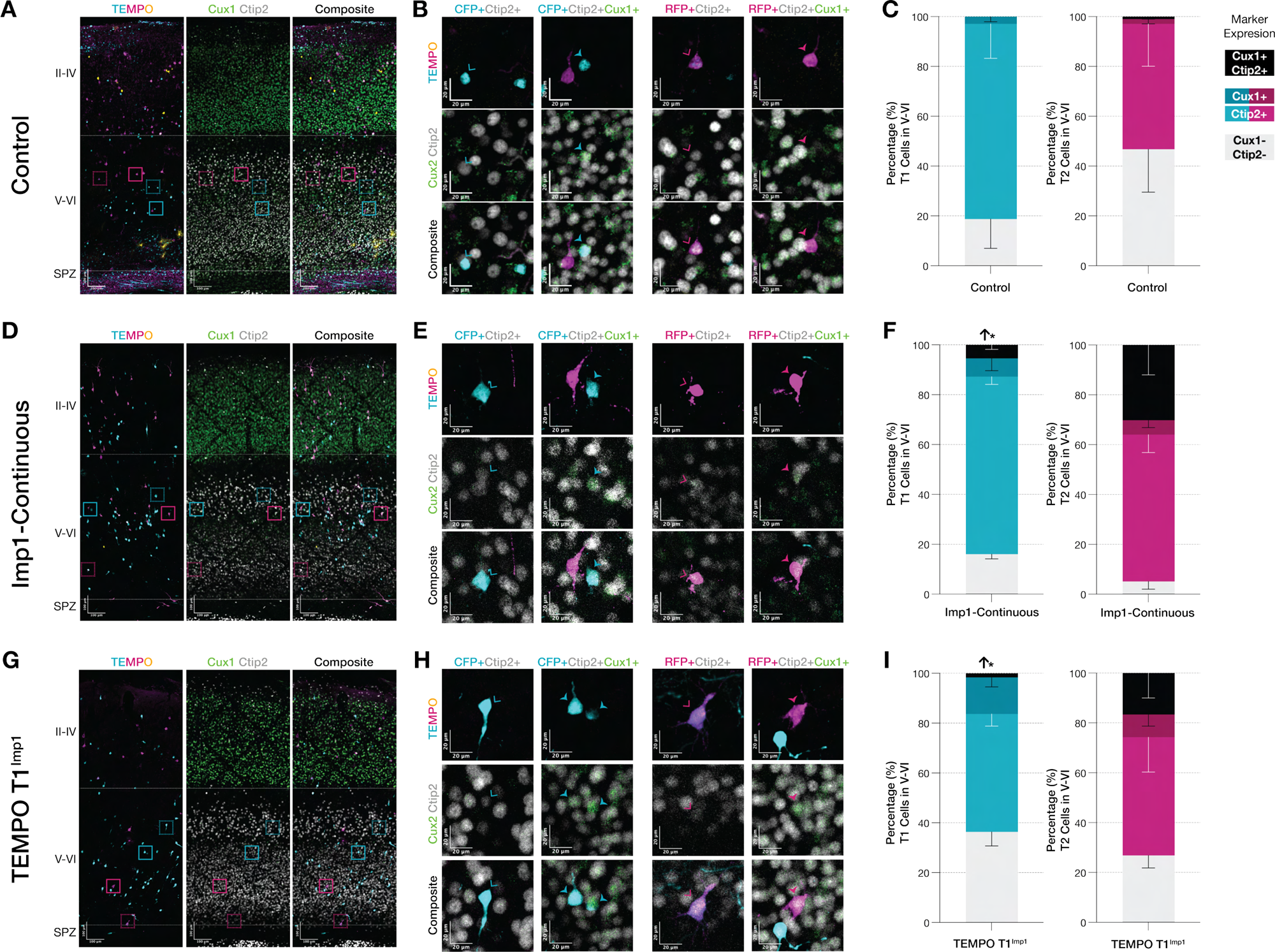
Imp1-mediated laminar redistribution reflects genuine changes in neuronal fate identity rather than migratory defects. **(A,D,G)** Representative images of control, continuous Imp1 overexpression, and T1^Imp1^ conditions showing TEMPO reporters, Cux1/Ctip2 immunostaining and overlays. Boxed regions: Ctip2+ TEMPO neurons (dashed) or double-positive Cux1+/Ctip2+ TEMPO cells (solid). **(B,E,H)** High magnification images of boxed regions highlight CFP-/RFP-labeled neurons in layers V-VI colocalizing with Ctip2 (outlined arrowheads) or double-positive for both markers (solid arrowheads). **(C,F,I)** Quantification of marker expression in CFP+ and RFP+ neurons residing in layers V-VI. Following continuous or T1^Imp1^ overexpression, neurons in deep layer maintain appropriate deep-layer molecular identities (predominantly Ctip2+), demonstrating that laminar distribution reflects bona fide fate specification changes rather than mislocalization. **(C)** In control conditions, CFP+: Ctip2 (78.31% ± 13.82%), Cux1 (2.93% ± 2.11%), double-negative (18.76% ± 11.73%). RFP+: Ctip2 (50.35% ± 17.01%), Cux1 (1.85% ± 1.85%), double-negative (46.76% ± 17.20%), double-positive (1.04% ± 1.04%). **(F)** Following continuous Imp1 overexpression, CFP+: Ctip2+ (71.18% ± 3.06%), Cux1+ (7.38% ± 4.93%), double-negative (16.06% ± 1.93%), double-positive (5.38% ± 1.80%). RFP+: Ctip2+ (58.92% ± 7.18%), Cux1+ (5.84% ± 3.01%), double-negative (5.10% ± 3.12%), double-positive (30.14% ± 11.98%). **(I)** In T1^Imp1^ overexpression, CFP+: Ctip2+ (47.26% ± 4.88%), Cux1+ (14.63% ± 3.76%), double-negative (36.41% ± 5.64%), double-positive (1.68% ± 0.57%). RFP+: Ctip2+ (47.38% ± 13.91%), Cux1+ (9.17% ± 4.68%), double-negative (26.86% ± 5.03%), double-positive (16.59% ± 10.02%). Dashed lines: upper (II-IV), lower cortical layers (V-VI) and subplate zone (SPZ). Scale bars: (A,D,G) 100 µm and (B,E,H) 20 µm. Data show mean±SEM. Statistics: two-tailed unpaired Welch’s t-test (*P < 0.05, **P < 0.01, *** P < 0.001).

Continuous Imp1 overexpression did not significantly alter the molecular identities of T1 neurons residing in lower layers (Figure 3D,E). The proportions of Ctip2+ (71%), Cux1+ (7%) and double-negative populations (16%) within the T1 cohort remained statistically unchanged compared to controls (Figure 3F, left). This finding strongly indicates that the CFP+ cells enriched in lower laminar layers following Imp1 induction represent bona fide deep-layer neurons rather than mislocalized upper-layer cells. Different from control brains, however, we observed around 5% of neurons co-expressing both Cux1 and Ctip2, suggesting that sustained early Imp1 expression may promote mixed or transitional fate specification in some early-born neurons. RFP+ neurons in layers V-VI following continuous Imp1 overexpression showed similar adherence to deep-layer identity, with marker distributions statistically comparable to controls: Ctip2+ (59%), Cux1+ (6%), and double-negative (5%) (Figure 3F, right), indicating that T2-born neurons redirected to deep layers also acquire appropriate deep-layer molecular identities. While an apparent increase in double-positive cells (30%) was observed in the RFP+ cohort compared to controls (1%), this difference did not reach statistical significance, likely due to the substantial variance in RFP+ measurements reflecting both the lower baseline numbers of T2-born neurons in deep layers and potentially greater fate heterogeneity. Nonetheless, the overall pattern suggests that both T1 and T2 neurons adopt molecular identities consistent with their laminar positions following Imp1 manipulation.

T1-restricted Imp1 overexpression produced nearly identical molecular patterns that parallel those observed with control brains (Figure 3G,H). T1^Imp1^ yielded statistically similar proportions of Ctip2+ neurons (47%), Cux1+ (15%), and double-negative populations (36%) in lower layers compared to controls, with a small percentage of double-positive cells (2%) also detected (Figure 3I, left). RFP+ neurons showed comparable patterns with Ctip2+ (47%), Cux1+ (9%), double-negative (27%), and double-positive (17%) populations (Figure 3I), though again with high variance reflecting smaller sample sizes. The high similarity of molecular identity profiles between T1^Imp1^ and continuous Imp1 overexpression demonstrates that early-stage Imp1 expression is sufficient to maintain progenitor competence for deep-layer neuron production and to redirect both T1 and subsequent T2 cohorts toward deep-layer fates. Across all conditions examined, the correspondence between laminar position and molecular identity establishes that Imp1-mediated redistribution reflects genuine changes in neuronal fate specification rather than migration defects, positioning Imp1 as a key regulator of temporal fate progression

### Live imaging reveals continuous cellular accumulation in T2^Imp1^ subplate regions

We next investigated the cellular basis behind the formation of ectopic subplate accumulations that showed distinct temporal window sensitivity. Prominent accumulations of RFP-labeled neurons in the subplate zone of P10 T2^Imp1^ brains suggested that Imp1 expression during this temporal window disrupts the normal radial migration of newborn neurons from the ventricular zone to their cortical destinations. To trace the developmental origins of these cells, we performed long-term live imaging of organotypic cortical slices prepared from E14.5 embryos previously electroporated at E12.5 with T2^Imp1^ TEMPO constructs. Slices were maintained *in vitro* and imaged every 30 minutes over 48-hours, enabling continuous observation of neuronal movements within defined germinal and migratory compartments (Figure 4A,B).

**Figure 4.**
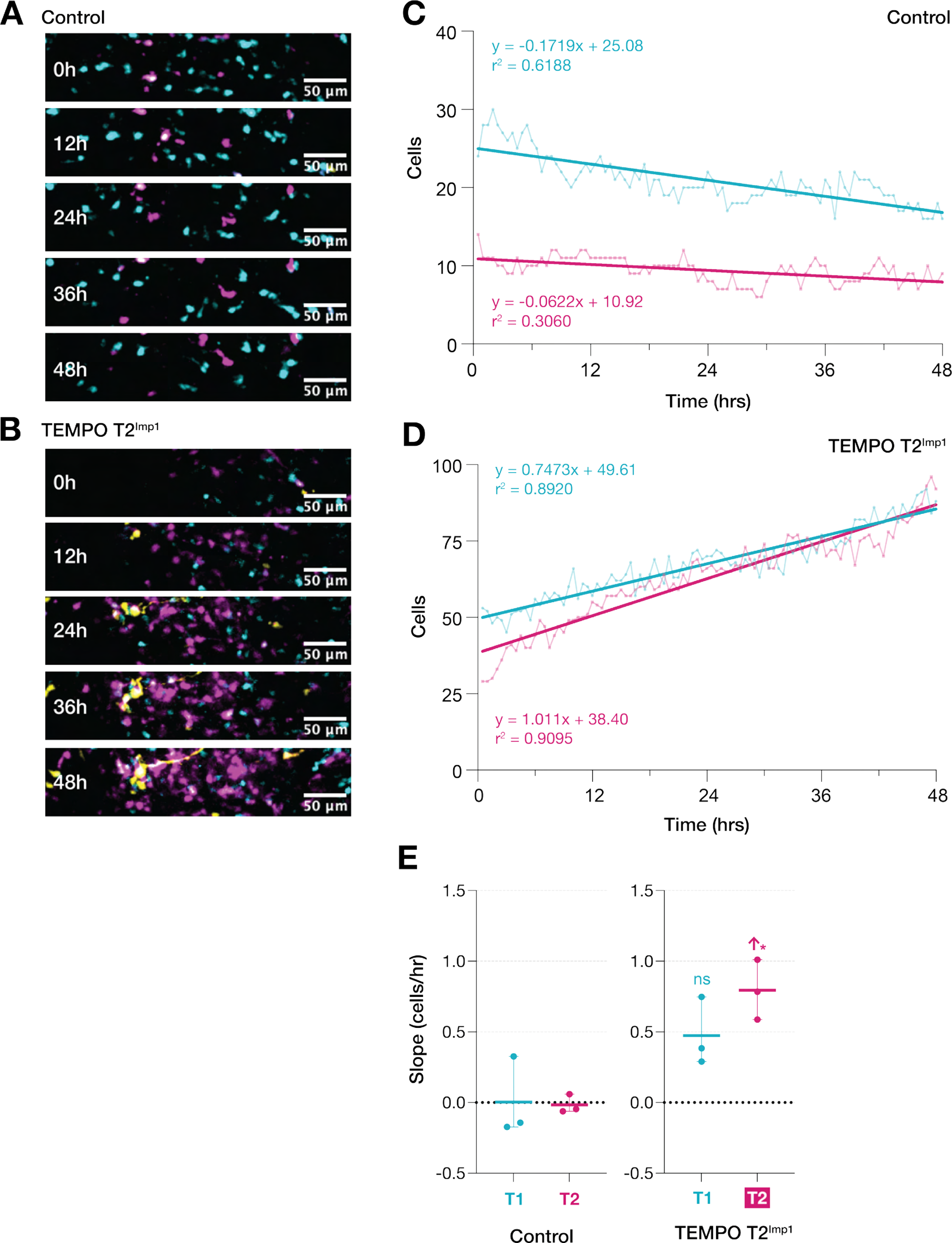
Continuous neuronal recruitment into the IZ-SVZ-VZ compartment underlies T2^Imp1^-induced RFP+ neuronal accumulation. (**A,B)** Representative time-lapse sequences (48 hours) from a single control and T2^Imp1^ brains. **(C)** In control, modest and variable changes in cell populations were observed (CFP+: slope = -0.172 cells/hr, r^2^ = 0.619; RFP+: slope = -0.062 cells/hr, r^2^ = 0.306). **(D)** T2^Imp1^ exhibits highly linear increases (CFP+: slope = 0.747 cells/hr, r^2^ = 0.892; RFP+: slope = 1.011 cells/hr, r^2^ = 0.910), suggesting continuous cellular recruitment. **(E)** Across brains (n = 3), RFP+ accumulation rate significantly increases in T2^Imp1^ (*P < 0.05). RFP+ accumulation exceeds CFP+, predicting eventual RFP+ dominance.

Given that the P10 accumulations localized immediately beneath the cortical plate, we reasoned that their embryonic precursors might be detectable in the intermediate zone (IZ), subventricular zone and VZ. Regions of interest (ROIs) were therefore delineated from the IZ, demarcated by prominent fiber tracts, through the SVZ to the VZ lining the lateral ventricle. Equivalent regions were selected in control slices to match size, anatomical position and baseline TEMPO+ cell density. Since YFP+ cells remained sparse at this developmental stage and did not form subplate accumulations in P10 brains, quantitative analyses were focused on CFP+ and RFP+ populations.

Time-lapse sequences revealed strikingly different population dynamics between conditions. In control slices, both CFP+ and RFP+ neurons exhibited modest and variable changes in number over time, reflecting a slightly biased outflow of cells from the ROI (Figure 4B). In contrast, T2^Imp1^ slices displayed steady, highly linear increases in both CFP+ and RFP+ populations (Figure 4D), with correlation coefficients exceeding 0.89 for both populations. This continuous, time-dependent accumulation indicates sustained cellular buildup within the IZ-SVZ-VZ compartment following T2^Imp1^ manipulation. Notably, the RFP+ accumulation rate consistently exceeded that of CFP+ cells (RFP+ slope = 1.011 cells/hr vs. CFP+ slope = 0.747 cells/hr), with this rate difference being statistically significant compared to controls (*P* < 0.05) (Figure 4E). This differential accumulation if sustained beyond the 48-hour imaging window, predicts that RFP+ neurons would eventually outnumber CFP+ neurons in this compartment.

The preferential enrichment of RFP+ neurons within this compartment during embryonic stages provides a developmental link to the RFP-dominant subplate accumulations observed at P10 (Figure 2C,D). The sustained accumulation dynamics, combined with the selective enrichment of RFP+ neurons specifically within the SPZ at postnatal stages (Figure 2B,E), establish that T2^Imp1^ expression alters the normal developmental progression through this region.

### T2-specific Imp1 induction triggers morphologically diverse glial foci with abnormal cellular organization independent of Hmga2

During analysis of P10 brain sections, we observed discrete glia-like accumulations scattered throughout the cortical parenchyma following T2-specific Imp1 overexpression and, to a lesser extent, continuous Imp1 conditions (Figure 5A). These aberrant foci were absent in controls and excluded from our quantification of normal glial development (Figures 1,2), as they represent aggregated focal assemblies rather than the spatially dispersed, glial populations that typify developmental gliogenesis (Bushong et al., 2004; Gallo & Deneen, 2014; Halassa et al., 2007; Nedergaard et al., 2003). In normal cortical development, individual radial glial clones typically generate small numbers of astrocytes (ranging from 2-50 cells) that disperse widely to tile the cortex, with mature astrocytes extending elaborate processes while minimizing overlap with neighboring cells (Beattie et al., 2017; Clavreul et al., 2019, 2022; García-Marqués & López-Mascaraque, 2013; Ojalvo-Sanz & López-Mascaraque, 2021).

**Figure 5.**
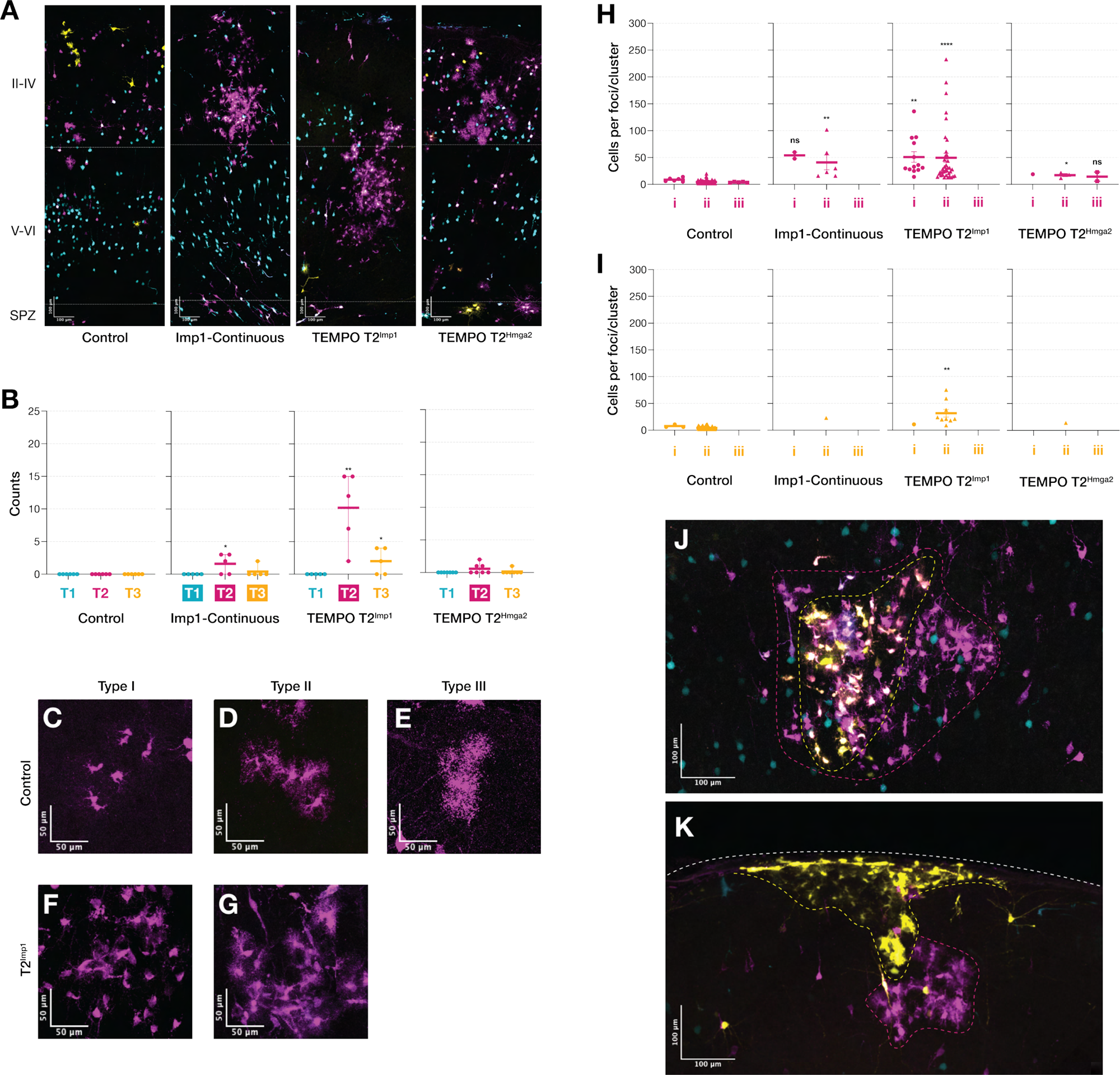
T2^Imp1^ induction triggers morphologically diverse glial foci with abnormal cellular organization through HMGA2-independent pathways. (**A**) Representative cortical sections showing RFP+ glial clusters at P10. (**B**) T2^Imp1^ produces significant RFP+ (10.20 ± 2.518 foci per brain) and to a lesser extent in continuous Imp1 overexpression (1.600 ± 0.6782 foci per brain), identifying T2 as a temporally sensitive window for Imp1-mediated glial clustering. T2^Imp1^ also shows YFP+ glial clusters (2.000 ± 0.8944 foci per brain). T2^Hmga2^ fails to induce significant glial clustering, demonstrating Imp1-specific mechanisms. (**C-H**) Morphological heterogeneity within glial clusters: **Type I** cells (minimal processes); **Type II** cells (characteristic astrocytic morphology); **Type III** cells (highly elaborate arbors). (**I**) RFP+ quantification shows dramatic expansion in T2^Imp1^. Glia in control brains organize into small Type II clusters (n = 105, ∼4 cells/cluster), sparse Type I (n = 6, ∼8 cells/cluster) and Type III (n = 5, ∼4 cells /cluster). Continuous Imp1 overexpression produces Type I (n = 2) and Type II foci (n = 6, ∼41 cells/focus, P < 0.05). T2^Imp1^ dramatically increases Type I (n = 13, ∼51 cells/focus, P < 0.001) and Type II foci (n = 38, ∼50 cells/focus, P < 0.0001) foci showing significant expansion. T2^Hmga2^ produces minimal foci across all types (Type I: n = 1; Type II: n = 3, ∼17 cells/focus, P < 0.05; Type III: n = 2), distinguishing Imp1 from Hmga2-mediated pathways. **(J)** YFP+ populations show similar RFP+ patterns with reduced frequency. Controls display Type I (n = 3, ∼8 cells/cluster) and Type II (n = 74, ∼4 cells/cluster). Continuous Imp1 and T2^Hmga2^ overexpression produce single YFP+ foci (∼23 and ∼14 cells per focus, respectively). T2^Imp1^ yields predominantly Type II foci (n = 9, ∼32 cells/focus, P < 0.05), consistent with the dramatic foci phenotype observed in RFP+ populations. **(K-L)** Spatial patterns for YFP+ foci: (K) nested YFP+ within RFP+ foci, and (L) layered arrangement with YFP+ near pial surface and RFP+ below. Dashed lines: upper (II-IV), lower (V-VI), SPZ. Scale bars: (A) 100 µm, (C-H) 50 µm. Data show mean ± SEM. Statistics: two-tailed unpaired Welch’s t-test (*P < 0.05, **P < 0.01, ***P < 0.001, ****P < 0.0001).

Quantitative analysis revealed striking enrichment of glial foci following T2^Imp1^ overexpression with RFP+ foci averaging at ∼10 per brain compared to ∼2 in continuous Imp1 conditions (Figure 5B). YFP+ foci (∼2 per brain) were also detected in T2^Imp1^ conditions, potentially reflecting molecular perdurance effects beyond the direct T2 window. To investigate whether Hmga2, a validated Imp1 downstream target promoting neural stem cell proliferation and self-renewal (Busch et al., 2016; Degrauwe et al., 2016; Nishino et al., 2008, 2013) mediated this phenotype, we performed T2-specific Hmga2 overexpression (T2^Hmga2^). This manipulation failed to produce significant glial foci (∼0.5 foci per brain; Figure 5B), indicating that foci formation operates via Imp1-specific mechanisms independent of this canonical self-renewal pathway.

Morphological examination revealed substantial heterogeneity within glial populations which we classified into three distinct categories based on process elaborations and arbor complexity (Figure 5C-G). Type I cells exhibited minimal process extension with compact somata resembling immature glia (Figure 5C, F). Type II cells showed characteristic astrocytic morphology with moderately branched processes forming territorial domains typical of mature cortical astrocytes (Figure 5D, G). Type III cells displayed highly elaborate arbors with extensive fine process ramification exceeding typical astrocytic complexity (Figure 5E).

Quantification of RFP+ populations revealed striking differences between conditions (Figure 5H). Control brains contained predominantly Type II clusters (n = 105, ∼4 cells/cluster), with sparse Type I (n = 6, ∼8 cells/cluster) and Type III (n = 5, ∼4 cells/cluster) representations. Continuous Imp1 overexpression produced Type I (n = 2) and Type II (n = 6, ∼41 cells/focus, P < 0.05). T2^Imp1^ dramatically increased both frequency and cellular composition, with Type I (n = 13, ∼51 cells per focus, P < 0.001) and Type II (n = 38, ∼50 cells/focus, P < 0.0001) foci showing significant expansion. Notably, Type III foci were completely absent from both Imp1 conditions. T2^Hmga2^ overexpression which produced minimal overall foci but included all three morphological types (Type I: n = 1; Type II: n = 3, ∼17 cells/focus, P < 0.05; Type III: n = 2), further distinguishing Imp1-specific effects from Hmga2-mediated pathways. YFP+ populations mirrored RFP+ patterns but with reduced overall frequency (Figure 5I). Control brains displayed Type I (n = 3, ∼8 cells/cluster) and Type II (n = 74, ∼4 cells/cluster) organizations. Continuous Imp1 and T2^Hmga2^ overexpression each produced minimal YFP+ foci (single Type II focus, ∼23 and ∼14 cells per focus, respectively) while T2^Imp1^ yielded predominantly Type II foci (n = 9, ∼32 cells/focus, P < 0.05).

Beyond the abnormal cellular expansion within individual foci, we observed stereotyped spatial organization patterns that further distinguished these structures from normal glial distributions. Examination of YFP+ Type II foci in T2^Imp1^ brains revealed distinct organizational motifs each representing approximately one-third of all YFP+ Type II foci (Figure 5J,K). First, YFP+ foci were contained within larger RFP+ foci, creating nested structures (Figure 5J). Second, YFP+ foci localized near the pial surface in layer I with adjacent RFP+ Type II foci positioned immediately below (Figure 5K). The remaining one-third consisted of YFP+ foci without clear spatial association with RFP+ populations. These stereotyped spatial arrangements combined with discrete spatial clustering, substantial cellular expansion (up to 12-fold greater than normal), and morphological heterogeneity suggest coordinated cellular behaviors rather than random aggregation, consistent with clonal expansion from individual Imp1-overexpressing progenitors, though single-cell lineage tracing would be required to confirm clonality.

## DISCUSSION

Corticogenesis unfolds as a temporally ordered program in which radial glia cycle through shifting competence states to produce deep-layer neurons, superficial-layer neurons and glia (Beattie & Hippenmeyer, 2017; Casingal et al., 2022; Nowakowski et al., 2016; Rakic, 1988). Within this framework, our data position Imp1 as a dosage- and window-dependent rheostat that paces fate transitions and when mistimed, disrupts neuronal migration and glial organization. Using TEMPO to co-register birth order with temporally restricted perturbations, we show that sustained Imp1 maintains early neurogenic programs, early (T1) pulses bias subsequent output, and later (T2) pulses unmask subplate and postnatal glial phenotypes absent from continuous or early-only manipulations. Through IUE, this mosaic gain-of-function approach builds upon previous conditional knockout studies (Nishino et al., 2013) by isolating cell-autonomous Imp1 functions within an otherwise normal cortical environment, revealing that observed phenotypes – including developmental arrest and altered laminar specification, reflect intrinsic regulatory capacity rather than secondary tissue-wide effects. We integrate these findings with broader themes in neurogenesis and outline a working model for how a post-transcriptional rheostat can orchestrate multi-scale cortical patterning.

### Imp1 operates as a competence preserving temporal regulator

Our findings reveal that Imp1 functions as a critical temporal gate determining when radial glial cells transition between developmental competence states, building directly upon foundational work by Nishino et al. (2013) demonstrating Imp1’s role in promoting fetal neural stem cell expansion while preventing premature differentiation. Their loss-of-function studies established that Imp1 deficiency accelerates neuronal and glial differentiation through HMGA2 stabilization and let-7 regulation, yet the temporal precision of Imp1’s regulatory control remained unclear from these constitutive manipulations.

Our temporal window-specific approach reveals Imp1 as a developmental rheostat whose effects depend critically on the expression timing. Continuous Imp1 expression indefinitely arrests RGCs in their early competence state, limiting them to producing deep-layer neurons and blocking gliogenesis. This suppression is evident when glial clusters are considered as clonal units, specifically continuous Imp1 brains contain far fewer glial clusters than controls (Figure 5I) suggesting dramatically reduced glial clone production. T1-restricted induction produces a partial arrest, temporarily holding RGCs at an earlier developmental state and biasing toward deep-layer specification with modest gliogenic delays, but permits progression to resume once the T1 window concludes. T2-restricted expression reveals increasingly constrained regulatory scope where neurons are redistributed toward deeper layers without affecting subsequent cohorts or altering the neurogenic-to-gliogenic switch, demonstrating that Imp1’s capacity to reprogram fate specification diminishes as development advances. This differential responsiveness wherein identical perturbations produce qualitatively distinct effects depending on developmental context establishes Imp1 as a context-dependent regulator whose competence-preserving function is shaped by the molecular landscape present at each stage.

To understand Imp1’s position within cortical temporal patterning networks, we compared its function to established temporal regulators that control developmental progression through distinct mechanisms. Prdm16 drives the transition from mid- to late-neurogenesis by limiting chromatin accessibility at early-program enhancers and promoting intermediate progenitor formation (Baizabal et al., 2018; L. He et al., 2021; Ray et al., 2022). Loss of Prdm16 causes persistence of deep-layer Ctip2+ corticofugal neurons into later stages when upper-layer Satb2+ neurons should predominate (L. He et al., 2021). Our T1-restricted Imp1 produces phenotypically similar outcomes, suggesting that Prdm16 and Imp1 may exert opposing regulatory functions. While Prdm16 normally advances temporal progression by promoting chromatin transitions that enable later fates, Imp1 appears to restrain progression by stabilizing early-stage transcripts. In contrast, Foxg1 appears to show complementary function to Imp1. Foxg1 restrains premature gliogenesis by suppressing FGF signaling though targets like Fgfr3, maintaining neurogenic competence until appropriate stages (Bose et al., 2025; Brancaccio et al., 2010; Hou et al., 2020). Foxg1 loss of function prematurely advances the neurogenic-to-gliogenic switch while our Imp1 oeverexpression delays or suppresses it, suggesting both may function as developmental brakes on the gliogenic transition through different molecular pathways. The degree of Imp1-mediated delay proved dosage- and time-dependent. T1-restricted expression produced modest delays that allowed eventual gliogenic progression, while continuous expression achieved near-complete suppression. This suggests that sustained early competence programs are largely incompatible with gliogenic progression. Together, these comparisons reveal that while transcriptional regulators like Prdm16 and Foxg1 define available cellular states and gate major transitions, post-transcriptional mechanisms like Imp1 provide an additional layer of control by modulating stage-specific transcript stability. Imp1 thus functions as a temporal coordinator whose dosage and timing determine whether developmental transitions are temporarily delayed or entirely blocked.

The mechanistic basis for this developmental arrest function aligns with emerging understanding of m^6^A-dependent transcript turnover during cortical development. Deleting the m^6^A-writer Mettl14 or Mettl3 in embryonic cortex prolongs radial glial cell cycles and extends neurogenesis into postnatal stages (Yen & Chen, 2021; Yoon et al., 2017), while loss of the m^6^A-reader Ythdf2 impairs neural mRNA clearance and severely decreases neurogenic output (Heck et al., 2020; M. Li et al., 2018). Importantly, m^6^A-modified transcripts exhibit significantly lower half-lives throughout neurogenesis (Serdar et al., 2025). These studies establish an mRNA-level timer that clears stage-inappropriate transcripts to drive temporal progression.

We hypothesize that Imp1 induction stabilizes early-competence transcripts normally degraded through m^6^A-guided turnover, effectively “freezing” radial glial cells in their initial competence state by preventing RNA-level transitions that drive progression. Critically, the m^6^A epitranscriptome itself changes dynamically in radial glial progenitors across developmental stages (Yen & Chen, 2021; Yoon et al., 2017). This temporal remodeling of the m^6^A landscape may explain Imp1’s stage-dependent effects - at each developmental window, Imp1 encounters a distinct complement of m^6^A-modified transcripts, determining whether perturbations manifest as comprehensive developmental arrest, partial fate biases, or constrained organizational defects.

### Stage-restricted manipulations reveal distinct developmental vulnerabilities in subplate organization and glial patterning

Beyond competence preservation effects, our temporal window-specific approach uncovered phenotypes that continuous expression paradigms would obscure. T2-specific manipulation (around E13.5 – E14.5) revealed two outcomes absent from other conditions: subplate accumulations and focal glial aggregates. These T2-specific phenotypes illuminate distinct developmental vulnerabilities in tissue organization beyond immediate fate specification.

The subplate accumulations represent a striking departure from effects on temporal fate progression. Live imaging revealed continuous accumulation of both CFP+ and RFP+ neurons within the IZ-SVZ-VZ during the T2 window. The concurrent accumulation of CFP+ cells, despite Imp1 expression being restricted to T2-born RFP+ neurons, suggests non-cell autonomous effects where T2^Imp1^ perturbations disrupt the local migratory environment or physically impede transit of earlier-born cohorts. Critically, RFP+ cells accumulated at significantly faster rates than CFP+ cells, indicating that T2-born neurons directly expressing Imp1 are more severely affected and could explain the RFP+ dominance of the subplate accumulation in the postnatal brain. Analysis of TEMPO reporter proportions in postnatal brains further supports this interpretation. When cortical layers are quantified without SPZ, TEMPO reporter proportions remain unchanged in T2^Imp1^ conditions compared to controls. Including SPZ in the analysis reveals significant RFP+ and its accumulation frequency positively correlating with RFP+ percentage (r^2^ = 0.75). This selective SPZ enrichment, combined with preserved cortical proportions, argues against global proliferative dysregulation and instead points to altered developmental transit or retention of T2-born neuronal cohorts. The biological basis likely involves Imp1’s roles in regulating β-actin mRNA localization and translation in growth cones (Farina et al., 2003; Ross et al., 1997; Triantopoulou & Vidaki, 2022), processes critical for the multipolar-to-bipolar transition required for radial migration through the subplate. (Agrawal & Welshhans, 2021; Ohtaka-Maruyama et al., 2018; Welshhans & Bassell, 2011; Xie et al., 2013). Perturbing these processes could slow subplate transit or impair developmental exit from germinal zones, creating kinetic bottlenecks that result in cellular accumulation (Boitard et al., 2015; La Fata et al., 2014; Ohtaka-Maruyama et al., 2018; Tabata & Nakajima, 2003).

T2^Imp1^ also fundamentally disrupted normal glial organization, producing more RFP+ foci than continuous Imp1 with dramatic cellular expansion compared to normal control clusters. YFP+ foci showed similar abnormal expansion, indicating that Imp1’s effects persist beyond the direct T2 manipulation window, potentially through molecular perdurance or establishment of heritable cellular states. Critically, T2^Hmga2^ failed to recapitulate this phenotype despite Hmga2 being a well-established Imp1 downstream target that promotes neural stem cell self-renewal (Busch et al., 2016; Nishino et al., 2013) revealing that glial foci formation operates through Imp1-specific regulatory pathways. The morphological heterogeneity within foci, ranging from cells with minimal processes to cells with highly elaborate arbors, suggests that Imp1 influences both proliferation and differentiation trajectories.

The stereotyped spatial organization of glial foci provides critical mechanistic insights into their formation. YFP+ foci exhibited two distinct spatial patterns, each representing approximately one-third of all YFP+ Type II foci, indicating programmed cellular behaviors rather than stochastic aggregation. The nested pattern, wherein smaller YFP+ foci reside within larger RFP+ foci is consistent with continued proliferation and delayed TEMPO color progression as accelerated cell cycle kinetics may outpace single-strand annealing (SSA) repair needed to transit to the next reporter (Espinosa-Medina et al., 2023). The layered pattern wherein YFP+ foci localize near the pial surface in layer I with adjacent RFP+ foci positioned below suggests directed migration of later-born progeny. This organization parallels aspects of normal astrocyte development where progenitors exhibit blood vessel-guided migration in superficial cortex (Tabata et al., 2022), layer I harbors specialized pial and subpial interlaminar astrocyte populations (Falcone et al., 2019), and clonal analysis demonstrates sibling astrocytes can form spatially separated clusters across layers (Clavreul et al., 2019; García-Marqués & López-Mascaraque, 2013; Lanjakornsiripan et al., 2018). The discrete spatial clustering, substantial cellular expansion, morphological heterogeneity, and stereotyped organizational patterns are consistent with clonal expansion from individual Imp1-overexpressing progenitors, though definitive lineage tracing would be required to confirm clonality.

The abnormal proliferative expansion and disrupted spatial organization observed in T2^Imp1^ glial foci raise intriguing questions about how developmental regulatory mechanisms can be co-opted when dysregulated. Imp1 is normally expressed during embryonic development but becomes silenced postnatally, with aberrant re-expression in multiple cancers driving tumor progression (Degrauwe et al., 2016; Hamilton et al., 2013; Zhu et al., 2023). Notably, Imp1 is overexpressed in glioblastoma and promotes glioma stem cell maintenance and aberrant proliferation (Yang et al., 2023). While our developmental model differs fundamentally from tumorigenesis, the observation that ectopic Imp1 expression during a specific developmental window can disrupt normal glial territorial organization and induce localized cellular aggregation suggests shared underlying mechanisms. Both contexts likely involve Imp1-mediated stabilization of transcripts regulating proliferation, spatial organization, and territorial spacing. This connection highlights how temporal regulatory networks that normally coordinate orderly cortical development might, when dysregulated outside their appropriate developmental context, inadvertently activate cellular programs that share features with pathological states.

Several limitations in this work warrant consideration. Our gain-of-function approach through mosaic *in utero* electroporation, while advantageous for studying cell-autonomous effects, leaves open questions about how Imp1 loss-of-function would affect temporal patterning with equivalent temporal resolution. Additionally, our population-level analyses cannot definitively determine whether SPZ accumulations and glial foci represent clonal expansions or aggregations of independently affected cells. Future studies employing single-cell lineage tracing combined with TEMPO would enable clonal-level analysis. Our molecular readout was also limited primarily to TEMPO reporters and selected fate markers, providing snapshots of identity specification but not comprehensive transcriptomic profiles. Spatial transcriptomics profiling TEMPO+ populations at multiple developmental stages would reveal the full spectrum of molecular changes and identify specific transcript targets whose stabilization drives the observed phenotypes. Although there are studies linking Hmga2 with tumorigenesis (Mansoori et al., 2021; Morishita et al., 2013; Zhang et al., 2018; Zhou et al., 2021), the specific Imp1-dependent pathways responsible for glial foci formation remain unknown. TEMPO could enable serial inductions to test candidate factors for rescuing normal development, providing mechanistic insights into the hierarchical organization of temporal fate programs.

## CONCLUSION

Our findings establish a new framework for understanding how post-transcriptional mechanisms integrate with transcriptional and epigenetic regulatory layers to control cortical temporal patterning. Rather than operating as isolated switches, these regulatory systems form an interconnected network where transcriptional programs define available cellular identities, epigenetic modifications gate developmental transitions, and post-transcriptional controllers like Imp1 fine-tune the pace and coordination of these processes. The temporal window-specific approach enabled by TEMPO demonstrates the critical importance of precise temporal control in developmental studies. Many traditional loss- and gain-of-function paradigms may obscure stage-specific regulatory functions, as demonstrated by our discovery that T2-restricted manipulation reveals migration and organizational phenotypes completely absent from continuous expression paradigms. This methodological advance suggests that systematic temporal dissection of developmental regulators could uncover previously hidden layers of biological complexity.

Ultimately, our work establishes post-transcriptional temporal control as a critical and previously underappreciated layer of cortical patterning. By demonstrating how a single RNA-binding protein can function as a developmental rheostat—modulating multiple aspects of neurogenesis through stage-specific effects on transcript stability and translation—we provide new insights into the molecular mechanisms that ensure proper brain assembly and the potential consequences when these systems go awry.

## ACKNOWLEDGEMENTS

We would like to thank the Janelia Shared Resource especially the Surgery and Histology Teams for their technical support. We also thank Jui-Chun Kao and Mindy Mackey for administrative support.

## FUNDING

National Institutes of Health **R01-NS134890** (T.L.)

Howard Hughes Medical Institute (T.L.)

American Heart Association **25PRE1354535** (R.A.A.)

## AUTHOR CONTRIBUTIONS

Conceptualization – D.F., I.E.-M., T.L.; Methodology – D.F., I.E.-M., R.A.A., T.L.; Investigation – D.F., I.E.-M., R.A.A., R.A., J.L-G., E.J., C.-P.Y.; Writing – Original Draft – R.A.A., T.L.; Writing – review and editing – R.A.A., D.F., I.E.-M., T.L.; Visualization – R.A.A.; Supervision – D.F., I.E.-M., T.L.; Project Administration – D.F., T.L., I.E.-M; Funding Acquisition – T.L.

